# Early Exposure to Polycyclic Aromatic Hydrocarbons (PAHs) and Cardiac Toxicity in a Species (*Xenopus laevis*) with Low Aryl Hydrocarbon Receptor (AHR) Responsiveness

**DOI:** 10.1101/301846

**Authors:** Madison C. Sestak, Julia A. Pinette, Caithlin M. Lamoureux, Susan L. Whittemore

## Abstract

Polycyclic aromatic hydrocarbons (PAHs) are ubiquitous, persistent environmental contaminants, of which 16 are EPA-designated priority pollutants. Cardiotoxicity is observed in fish with developmental exposures to certain PAHs; however, the mechanism of toxicity can differ. Phenanthrene (PHE) and benzo(a)pyrene (BaP) are both cardiotoxic to fish, but PHE acts independently of aryl hydrocarbon receptor (AHR) activation while BaP-associated cardiotoxicity is AHR-dependent. To further understanding of mechanisms of toxicity, we compared the effects of early exposure to the priority PAHs pyrene (PYR), fluoranthene (FLA), PHE and BaP on cardiac function and *cytochrome P450 type 1A* (*cyp1a*) mRNA expression, an indicator of AHR activation, in a model system with lower AHR sensitivity than that of fish, the embryos and larvae of *Xenopus laevis.* Exposure to PYR, PHE, and FLA (0.25 – 25 μM) caused ventricular tachycardia early in heart development, but bradycardia and atrioventricular (AV) block in later stages. Elevated *cyp1a* mRNA levels indicate that FLA and BaP, but not PHE or PYR, are AHR agonists. The finding of FLA-induced cardiotoxicity and *cyp1a* expression (35-fold) is particularly surprising as FLA inhibits CYP1A activity in fish and, as a single compound, is not cardiotoxic. Our results suggest that early exposure to PHE, PYR, and FLA, but not to BaP, compromises cardiac function by altering normal pacemaker activity and conduction in *Xenopus*, effects associated with increased mortality. Our findings also reveal a considerable degree of species specificity between fish and frog regarding cardiac sensitivity to developmental PAH exposures and have implications for the cardiovascular health of PAH-exposed humans and wild amphibians.

## Introduction

Polycyclic aromatic hydrocarbons (PAHs) are a large group of ubiquitous and persistent pollutants that continue to be deposited into the environment primarily with the incomplete combustion of fossil fuels among other sources (Agency for Toxic Substances and Disease Registry (ATSDR), 1995). The Environmental Protection Agency (EPA) has designated 16 PAHs as priority pollutants. These include the four PAHs under investigation here: the tricyclic AH phenanthrene (PHE), the tetracyclic AHs pyrene (PYR) and fluoranthene (FLA), and the five-ringed benzo-(a)-pyrene (BaP) (Table 1). In contrast to BaP, a well-studied carcinogen, PHE, PYR and FLA toxicity remains poorly characterized and their inclusion as priority pollutants is primarily due to their prevalence in the environment.

**Table 1.** Dose range and molecular weights for PAHs

| PAH | Molecular Weight<br>g/mol | Dose Range<br>in $\mu\text{M}$ | Dose Range<br>in ppm | Dose Range<br>in $\mu\text{g/L}$ |
| --- | --- | --- | --- | --- |
| Phenanthrene (PHE) | 178.2 | 0.25 - 25 | 0.044 - 4.4 | 44 - 4400 |
| Pyrene (PYR) | 202.3 | 0.25 - 25 | 0.05 – 5 | 50 - 5000 |
| Fluoranthene (FLA) | 202.3 | 0.25 - 25 | 0.05 – 5 | 50 - 5000 |
| Benzo-(a)-pyrene (BaP) | 252.3 | 0.5 - 5 | 0.13 – 1.3 | 130 - 13000 |

Crude oil spills, most notably the Deepwater Horizon disaster in the Gulf of Mexico (2010), generated multiple investigations into the impact of PAH exposure on developing fish and fish populations and the resulting economic implications. It is now well-established that early exposure to select polycyclic aromatic hydrocarbons (PAHs), either as single compounds or as components of weathered crude oil, leads to abnormal cardiac morphology and function in fish. These studies also revealed diversity in modes of action and tissue specificity for single PAHs. For example, in zebrafish (*Danio rerio*), cardiotoxicity associated with developmental exposure to PHE is independent of the AHR (aryl hydrocarbon receptor) and CYP1A (cytochrome P450 type 1A) activity (Incardona et al., 2005), while BaP-induced cardiotoxicity is both AHR and CYP1A-dependent (Huang et al., 2012; Incardona et al., 2011). While there is little cardiotoxicity associated with PYR exposure, it is reported to cause AHR- and CYP1A-dependent perturbations in the form of anemia, peripheral vascular defects, and neuronal cell death in the early life stages of zebrafish (Incardona et al., 2004; Zhang et al., 2012). In contrast, FLA is an inhibitor of CYP1A activity in fish and, as a single compound, yet to be associated with abnormal cardiac function or morphology. Collectively, these studies underscore the diversity in target tissues and mechanism of toxicity among individual PAHs.

The embryos and larvae of the African clawed frog *Xenopus laevis* are important models for investigating normal vertebrate heart development, providing insight, for example, into human congenital heart defects (Kolker et al., 2000; Mohun et al., 2000; Warkman and Krieg, 2007). *Xenopus* also serves as a tractable model amphibian for identifying potential environmental impacts on amphibian species in general, an important attribute in the face of global decline in amphibian populations (Adams et al., 2013; Grant et al., 2016). Finally, *Xenopus* embryos and larvae provide an attractive system for investigating mechanisms of developmental toxicity with PAH exposure because, in comparison to fish and even rodent models, the sensitivity of their AHR system to certain AHR agonists (e.g. dioxin) is greatly reduced and more closely mirrors that of humans and other amphibians (Lavine et al., 2005; Silkworth et al., 2005; Zimmermann et al., 2008).

Humans are typically exposed to PHE, PYR, and FLA and other PAHs through the ingestion of contaminated food and the inhalation of polluted air (Marti-Cid et al., 2008; Narváez et al., 2008). Other factors include proximity to industrial and urban settings, smoking and work-related exposure (Agency for Toxic Substances and Disease Registry (ATSDR), 1995; Jung et al., 2010; Mahler et al., 2012; Williams et al., 2012). In addition, the use of biomass cookstoves in poorly ventilated shelters represents a significant route of exposure for a large proportion of the world population (Gadi et al., 2012; Riojas-Rodriguez et al., 2011).

For a variety of reasons, children bear a higher PAH burden than do adults (Perera, 2008). Furthermore, there is evidence for significant placental transfer of PHE, PYR, and FLA in humans. Indeed, PHE, PYR, and FLA are among the top five most abundant priority PAHs detected in human placenta, cord blood and milk, underscoring the need to investigate their potential for developmental risk (Tsang et al., 2011; Yu et al., 2011). In contrast, BaP was undetectable in human milk, maternal and cord serum in one of these studies (Tsang et al., 2011) and present at much lower levels than PHE, PYR and FLA in the other.

Wild larval amphibians are exposed to PAHs through contamination of their aquatic environments by oil spills, contaminated sediments, and with proximity to roadways and industrial settings (Bradford et al., 2010; Garrigues et al., 2004). In particular, runoff from roadways and other paved surfaces that have been sealed with coal tar-based sealants represents a significant PAH source for watersheds particularly as the road or parking lot surface is degraded over time with vehicular use and weather conditions (Mahler et al., 2012). It should be noted that coal tar sealant dust has likewise emerged as a concerning source of PAH exposure in children particularly in urban areas (Williams et al., 2012).

There are few additional studies that have focused on the developmental effects of exposure to single PAHs to corroborate their distinct modes of action and tissue- or species-specificity. Understanding the selective developmental impacts and the underlying mechanisms by which different individual PAHs adversely alter development is critical to the accurate assessment of the risk of exposure to both humans and wildlife. To extend current understanding of the mechanism(s) of toxicity for these prevalent priority PAHs, we compared the impact of early exposure to the common pollutants PHE, PYR, FLA, and BaP on cardiac function and *cyp1a* mRNA expression using a model system in which sensitivity to AHR agonists is particularly low, providing a useful contrast to findings derived from fish studies. We predicted that, in comparison to fish, *Xenopus* hearts would be equally sensitive to PHE since its actions are AHR- and CYP1A-independent, but far less sensitive to BaP as its cardiotoxic effects are mediated by the AHR and CYP1A. In addition, we predicted that it would be unlikely that early exposure to FLA, a CYP1A inhibitor, or to PYR, only a moderate AHR agonist, would result in any negative impacts on cardiac function in *Xenopus.*

## Materials and Methods

*Animal husbandry and embryo manipulation* Albino and pigmented adult *Xenopus laevis* (Nasco, Fort Atkinson, WI) were maintained in our frog facility under a 12 h:12 h light:dark cycle at 19°C. We used albino frogs to assess cardiac function and pigmented frogs for all other studies. Prior to breeding, males and females were injected with human chorionic gonadotropin (Sigma Aldrich) in the dorsal lymph sac to induce breeding and placed overnight in breeding chambers containing 10% Holtfreter’s solution (HF, saline). Embryos were dejellied in 2% cysteine (in 10% HF, pH 8.0) for 5-7 min and then maintained in 10% HF until the onset of gastrulation or Nieuwkoop and Faber (NF) stage 10.5 (Nieuwkoop and Faber, 1967). All animals were treated humanely according to approved Institutional Animal Care and Use Committee protocols.

*Embryo exposure protocol* We chose a range of doses similar to those used in several fish studies (0.25 - 25 μM for PHE, FLA, and PYR and 0.5 – 5 μM for BaP) to facilitate comparison and determination of species-specific effects (see Table 1 for doses in ppm). Stock solutions of fluoranthene, benzo(a)pyrene, pyrene, and phenanthrene (>96-99% purity, Sigma Aldrich) were prepared in 100% dimethyl sulfoxide (DMSO, Sigma Aldrich) and stored at -20 °C. All other reagents were obtained from Sigma Aldrich unless otherwise noted. Treatments consisted of dilutions of the stocks with 10% HF to achieve final concentrations. The experiment was replicated using 6 different broods of embryos (resulting from 6 different matings) and included age-matched, untreated, vehicle (0.02-0.04% DMSO) controls. We previously established that vehicle controls did not differ from 10% HF controls. 10 embryos from each brood were randomly assigned to the vehicle control or one of the 11 PAH treatment groups and housed in glass scintillation vials at a density of 1.5 mL/embryo at 22°C in 12L:12D. Observations were recorded and media changed daily. If allowed to develop beyond NF stage 45 (the onset of feeding), larvae were fed finely ground tadpole brittle (Nasco, Fort Atkinson, WI). Embryos and larvae were anesthetized where indicated and euthanized with tricaine (MS-222) at completion of experiment.

*Mortality*, *Morphometric*, *and Edema Assessment* Embryos from the same brood were exposed continuously to 0.25, 2.5 or 25 μM PHE, PYR or FLA or to 0.5 or 5 μM BaP from day 0 (NF stage 10.5) to day 10 (typically NF stage 48-49). Media changes and monitoring for mortality occurred daily. Snout-to-vent lengths were obtained for embryos exposed for 96 h to 0.25, 2.5 or 25 μM PHE, PYR or FLA or to 0.5 or 5 μM BaP after fixation for 1 h in MEMFA (100 mM MOPS, pH 7.4; 2 mM EGTA; 1 mM MgSO_4_; 4% paraformaldehyde), dehydrated in methanol, and stored at -20 °C. Four different broods were used for these experiments. 12 embryos/brood were measured and examined for craniofacial, skeletal, and eye abnormalities and edema of the pericardium and gut by an observer blinded to the treatment group.

*Heart rate and atrioventicular block* Animals were exposed to either 0.25, 2.5 or 25 μM PHE, PYR or FLA or to 0.5 or 5 μM BaP for the time periods indicated below and their heart rates (HR) and incidence of atrioventricular (AV) block compared to animals exposed to 0.02% DMSO (vehicle or negative control). HRs were recorded at NF stage 37 (48 h exposure, post-atrioventricular partitioning), NF stage 42 (72 h exposure, during period of valve formation), NF stage 45 (96 h exposure, during atrial partitioning), and NF stage 46/47 (120 h exposure, mature heart phenotype) (Nieuwkoop and Faber, 1967). Embryos were allowed to equilibrate to room temperature for 60 m prior to video recording. Animals at stages 45 and 46/47 were first anesthetized with MS-222, at a dose and time period previously established to not alter HR, and positioned such that both the ventricular and atrial HR could be determined. Only ventricular HRs were obtained for the earlier stages.

Video recordings of beating hearts were captured using a Leica™ camcorder mounted to a dissecting microscope and Leica™ video capture software. Videos were analyzed using Camtasia™ video-editing and Image J™ (NIH) software to determine HR (in beats per minute, bpm) for a 20 s period. For the latter two NF stages, the ratio of atrial to ventricular contractions for the 20 s period was determined to assess for the presence and extent of AV block. For all of the above measures, the observers were blinded to the treatment group until measurements were complete.

*qRT*-*PCR for cyp1a mRNA levels* For qRT-PCR analyses, mRNA was purified using RNaqueous™ kit followed by TURBO DNA-free™ treatment (both Life Technologies, Carlsbad CA, USA) from NF stage 42 whole embryos exposed to PHE, PYR, FLA at 0.25, 2.5, 25 μM, B(a)P at 5 μM, or 0.02% DMSO for 72 h. Each sample (n=12) represented a pool of 7-9 embryos obtained from three different matings. RNA was quantified by measuring absorbance at 260 nm (Nanodrop™) and stored at -20 °C. RNA was reverse transcribed using a High Capacity RNA to cDNA™ kit (Life Technologies) and quantitative RT-PCR was performed using a StepOnePlus™ System (Applied Biosystems, Inc., ABI) and Taqman^®^ Gene Expression Assays (ABI) for *cytochrome P450 type 1a6* (*cyp1a6*), *cytochrome P450 type 1a7* (*cyp1a7)* (Fujita, 1999), and *ornithine decarboxylase (odc1*, endogenous control) mRNA (Sindelka et al., 2006). As a pseudotetraploid, *Xenopus laevis* possesses two *cyp1a* genes (Lavine et al., 2005). Primer and probe sequences are provided in Table 2. Samples were analyzed without reverse transcriptase as a control for genomic DNA contamination and without template DNA to control for reagent contamination. All gene expression assays were demonstrated to amplify a single product of expected size and with equal efficiency over a range of RNA concentrations (Bustin et al., 2009).

**Table 2.**
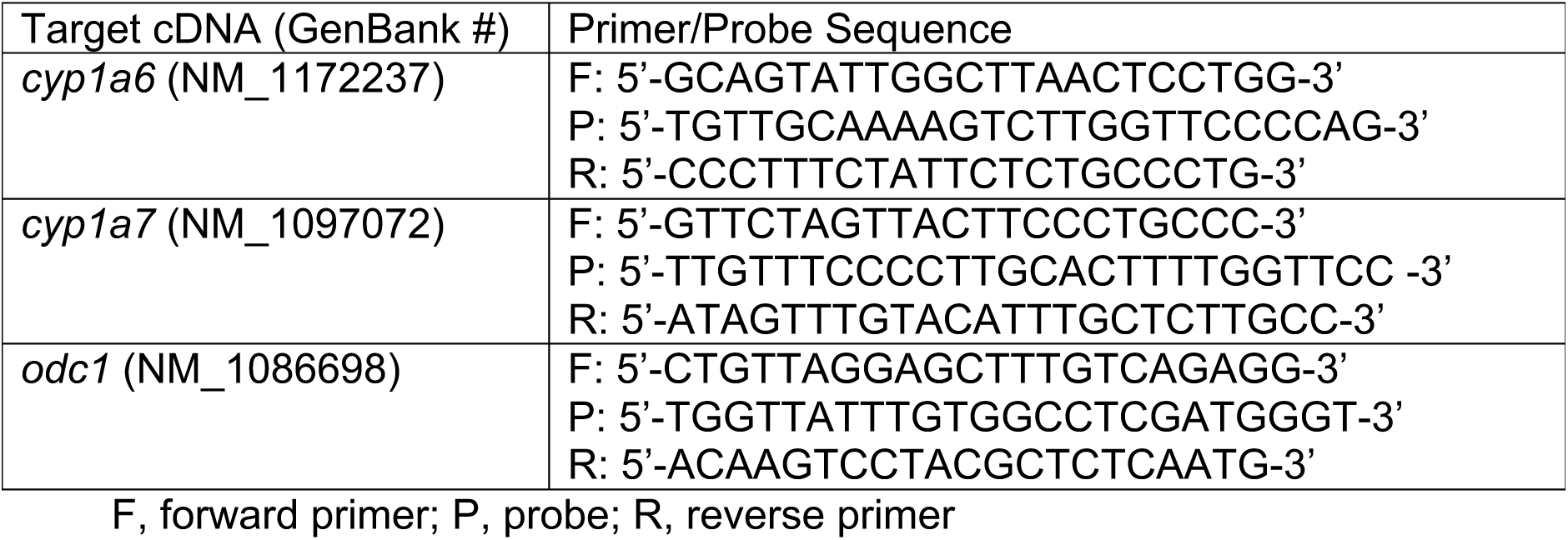
Primer and probe sequences used for quantitative RT-PCR experiments

*Statistics* Statistical significance (p ≤ 0.05) was determined using one-way analysis of variance (ANOVA) followed by Dunnett’s multiple comparisons test (GraphPad™ Prism 6.0, GraphPad Software, San Diego). The results of the gene expression studies were analyzed using the 2^-ΔΔCT^ method (Livak and Schmittgen, 2001) and expressed as fold increases over vehicle (0.02% DMSO) control mRNA levels.

## Results

*Mortality and Developmental Abnormalities* Overall, few developmental abnormalities were noted following 96 h PAH treatments (Table 3) and those few were typically mild, not severe, in nature. The most common abnormality, craniofacial defects, occurred in 7% of embryos exposed to B(a)P (0.5 and 5 μM) and FLA (25 μM). Edema was rarely observed. There was only a single observation of pericardial edema (with 0.25 μM PYR).

**Table 3.** Percentage of animals exhibiting developmental abnormalities and edema

| PAH | PYR<br>μM |  |  | PHE<br>μM |  |  | FLA<br>μM |  |  | BaP<br>μM |  |
| --- | --- | --- | --- | --- | --- | --- | --- | --- | --- | --- | --- |
|  | .25 | 2.5 | 25 | .25 | 2.5 | 25 | .25 | 2.5 | 25 | .5 | 5 |
| *Abnormalities:* |  |  |  |  |  |  |  |  |  |  |  |
| craniofacial | 4 | 4 | 2 | - | - | 2 | 2 | 4 | 7 | 7 | 7 |
| dorsal curvature | - | 2 | 2 | - | - | 2 | 2 | - | 4 | - | 2 |
| eye | 2 | - | - | - | - | - | - | 2 | - | 4 | - |
| *Edema:* |  |  |  |  |  |  |  |  |  |  |  |
| general | - | 2 | - | - | - | 2 | 2 | - | - | - | - |
| pericardial | 2 | - | - | - | - | - | - | - | - | - | - |

n = 45/treatment/6 batches; 96 h PAH exposure; no abnormalities or edema observed for DMSO controls

There was no evidence of developmental delay or advancement with PAH exposure; however, embryos exposed to 0.25 and 2.5 μM PHE had greater snout-to-vent lengths (p< 0.05) than those of vehicle controls (data not shown). No other differences in snout-to-vent lengths were observed. Finally, it was noted that animals exposed to 25 μM PHE, FLA, and PYR were overall much lighter in appearance with punctate, not dendritic, ventral melanophores.

Survival rates did not differ between PAH-treated and vehicle control animals at day 5 of exposure, but by day 8, the majority of animals exposed continuously to 2.5 and 25 μM PYR, 25 μM PHE and 25 μM FLA had died (Figure 1). This pattern remained until the end of the experiment at day 10. In contrast, the survival rates of animals exposed to the lower doses of FLA and PHE and to BaP did not differ from controls over the 10-day period.

**Figure 1.**
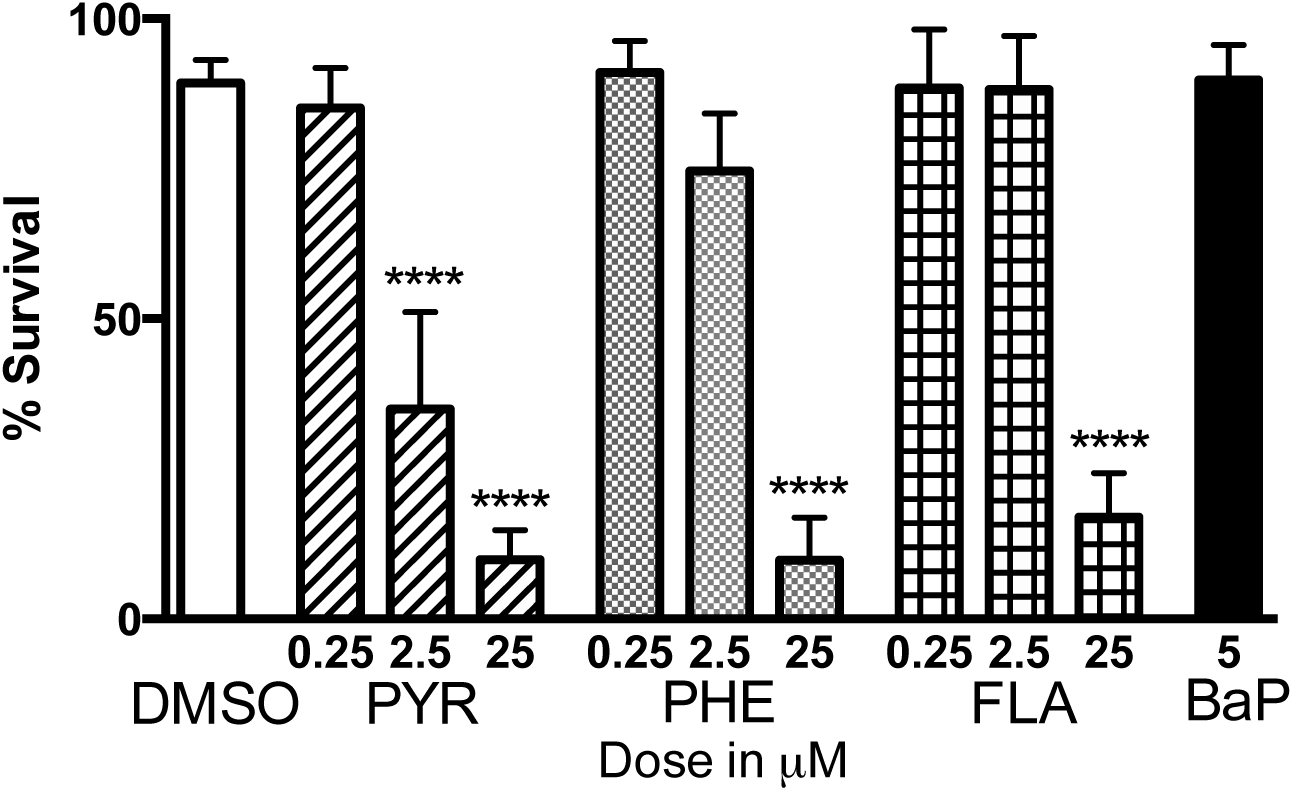
Percent survival for larvae following continuous exposure to PAHs or to 0.04% DMSO (vehicle control) for 8 days. Means ± S.E.M for 30 animals/batch with 6 batches/treatment. Data were analyzed using one-way ANOVA and Dunnett’s multiple comparisons tests; ^⋆⋆⋆⋆^ represents a difference (p ≤ 0.0001) from vehicle control. PYR = pyrene, PHE = phenanthrene, FLA = fluoranthene (at 0.25, 2.5 or 25 μM), and BaP = benzo(a)pyrene (at 5 μM).

*Heart rate* Ventricular rates for *X. laevis* embryos increased steadily with heart development (Figure 2), from a mean of 57 bpm for NF stage 37 DMSO controls (day 2, 48 hpe), immediately following cardiac looping and atrioventricular partitioning, to a high of 117 bpm for NF stage 46/47 animals (day 5, 120 hpe) with a mature heart phenotype. In NF stage 37 embryos, ventricular tachycardia (elevated HR) was observed with the highest dose of FLA when compared to vehicle controls. By NF stage 42 (day 3, 72 hpe, during heart valve formation), all PAHs at all doses, with the exception of 0.5 μM B(a)P, caused ventricular tachycardia.

**Figure 2.**
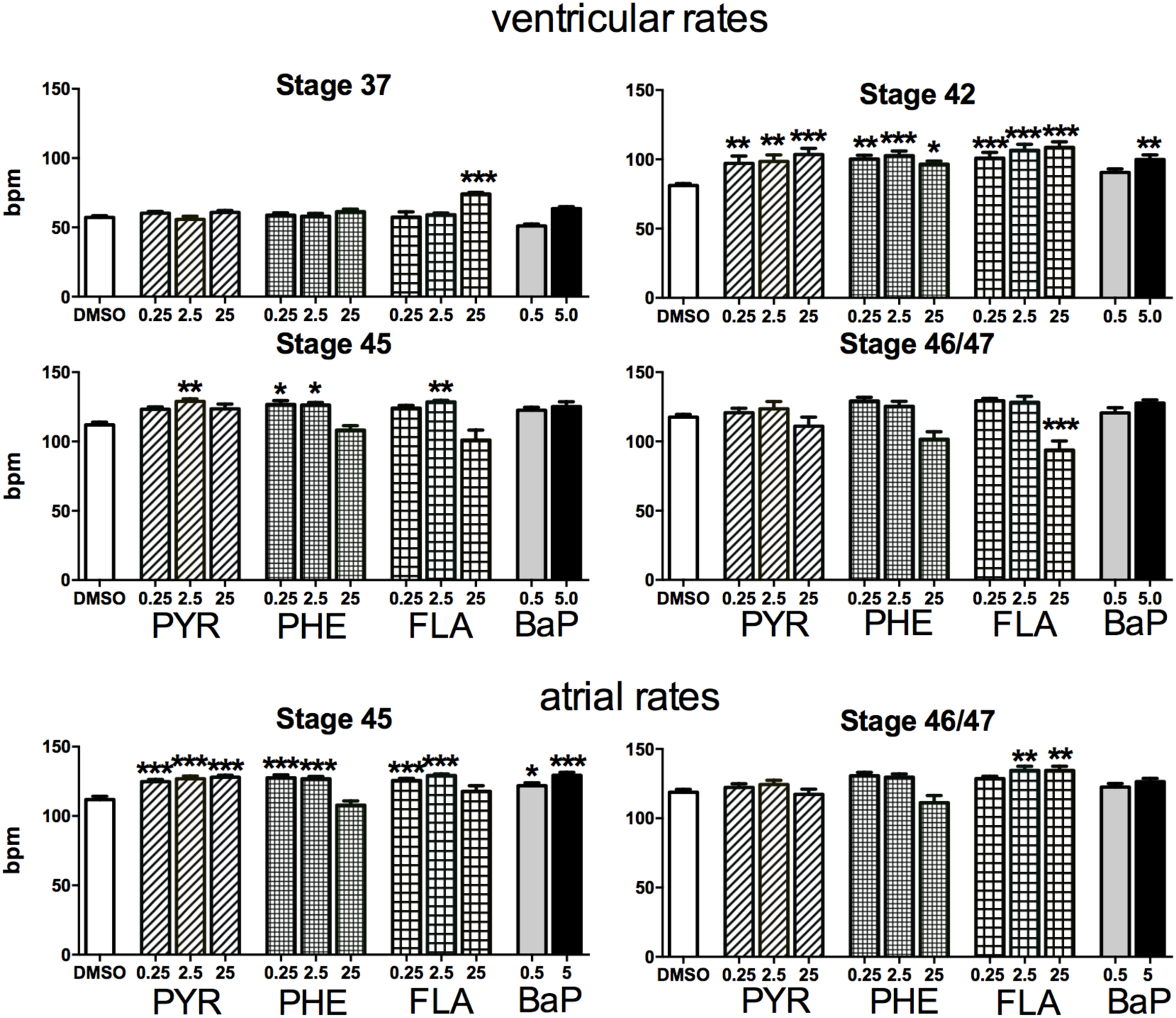
Ventricular rates in beats/min for vehicle controls (0.02% DMSO) and PAH-treated animals at NF stages 37 (day 2), 42 (day 3), 45 (day 4), and 46/47 (day 5). Atrial rates were obtained for stages 45 and 46/47 only. Means ± S.E.M for n=20-53 animals/treatment (6 batches); asterisks indicate a difference from the vehicle control (^⋆^ p <0.05, ^⋆⋆^ p <0.01, ^⋆⋆⋆^ p <0.001). PYR = pyrene, PHE = phenanthrene, FLA = fluoranthene (at 0.25, 2.5 or 25 μM), and BaP = benzo(a)pyrene (at 0.5 and 5 μM).

Ventricular tachycardia persisted into NF stage 45 (day 4, 96 hpe, during atrial partitioning) for many of the PAH-exposed embryos with the exception of those exposed to the highest dose (25 μM) of PYR, PHE and FLA where HR did not differ from those of vehicle controls. There were few differences between vehicle controls and animals at NF stage 46/47 (day 5, 120 hpe, mature heart phenotype). Ventricular bradycardia (decreased HR) was observed in larvae exposed to 25 μM FLA and while no other significant differences in ventricular HR were observed at this stage, there was a trend toward bradycardia for animals exposed to 25 μM PYR and PHE.

At NF Stage 45 (96 hpe), atrial HR for PAH-treated animals typically paralleled ventricular HR exhibiting tachycardia (Figure 2). By NF stage 46/47 (120 hpe), atrial HR for the treatment groups did not differ from vehicle controls. However, while exposure to either 25 μM FLA or PHE was associated, albeit to varying degree, with ventricular bradycardia, their effects on atrial HR differed. With PHE exposure, atrial and ventricular HR tended to mirror each other, both trending lower than control values, while in FLA-treated animals, atrial HR was higher and ventricular HR lower than those of control animals. In support of this difference, we did observe with PHE, but not with FLA treatment, several instances of AV delay, where a one: one correspondence between atrial and ventricular contractions was maintained, but the time between the atrial and ventricular contractions (A:V interval) was obviously increased (data not shown).

### Atrioventricular Block

Increased incidence of atrioventricular (AV) block, where the number of atrial contractions exceeded the number of ventricular contractions for a 20 s period (Table 4), was typically observed with PAH exposure and never observed for vehicle controls or with 0.5 μM B(a)P. The highest incidence (45% at stage 46/47) was noted for animals exposed to 25 μM FLA. Complete AV block (“ventricular standstill”) for the full 20 s period was observed with FLA, PYR and PHE exposure, but never with exposure to B(a)P.

**Table 4.** Incidence of atrioventricular (AV) block and full block (ventricular standstill) as percentage of PAH-exposed animals

|  | PYR<br>μM |  |  | PHE<br>μM |  |  | FLA<br>μM |  |  | BaP<br>μM |  |
| --- | --- | --- | --- | --- | --- | --- | --- | --- | --- | --- | --- |
|  | .25 | 2.5 | 25 | .25 | 2.5 | 25 | .25 | 2.5 | 25 | .5 | 5 |
| *Stage 45 (day 4)* |  |  |  |  |  |  |  |  |  |  |  |
| % AV Block | - | 7 | 19 | - | 13 | 11 | 3 | - | 36 | - | 7 |
| % Full Block | - | - | 3 | - | - | - | 3 | - | 11 | - | - |
| *Stage 46/47(day 5)* |  |  |  |  |  |  |  |  |  |  |  |
| % AV Block | 6 | 10 | 19 | - | 4 | 23 | - | 14 | 45 | - | 3 |
| % Full Block | - | 3 | - | - | - | 4 | - | - | 3 | - | - |

AV block = # atrial > # ventricular contractions and full block = no ventricular contractions for 20 sec (n = 24-34 animals/6 batches). PYR = pyrene, PHE = phenanthrene, FLA = fluoranthene, BaP = benzo(a)pyrene. No AV block for DMSO controls.

### cyplA mRNA levels

Changes in mRNA levels for the two *cyp1a* genes (*cyp1a6* and *cyp1a7*), following 72 h PAH treatment, were expressed as fold differences from DMSO control levels. In general, the *cyp1a6* paralog was less sensitive to PAH induction than was *cyp1a7*, with the exception of the highest dose of PHE (25 μM) which was associated with a 15-fold increase in *cyp1a6*, but no effect on *cyp1a7*, expression. As expected, the classic AHR agonist BaP (5 μM) was a potent inducer of *cyp1a* (~60-fold for *cyp1a7*) expression. Surprisingly, FLA, an inhibitor of *cyp1a* expression in zebrafish, was found to be an AHR activator (at 25 μM) in *Xenopus* embryos, as evidenced by a 35-fold increase in *cyp1a7* expression.

## Discussion

As the first organ to develop and due to its critical role in establishing the circulation, the heart serves as an important endpoint of developmental toxicity. Most, if not all, previous studies investigating the cardiotoxic potential of single PAHs used fish models. For two reasons, we chose to study PAH-induced cardiotoxicity in *Xenopus laevis* tadpoles. *Xenopus* hearts have three, not two, chambers and are more similar in structure and function to human hearts. The embryos and larvae of the African clawed frog *Xenopus laevis* have long served as important models for investigating normal vertebrate heart development (Kolker et al., 2000; Mohun et al., 2000; Warkman and Krieg, 2007).

Furthermore, fish and frogs differ in their relative sensitivities to contaminants that exert their toxicity through the binding and activation of the aryl hydrocarbon receptor (AHR). In comparison to frogs and other vertebrate models, fish possess a more complex AHR system. Due to gene and genome duplication events, fish possess two AHR clades, AHR1 and AHR2, and three AHR genes (*ahr1a*, *ahr1b*, and *ahr2*) (Hahn ME et al., 2006; Incardona et al., 2011). Importantly, the sensitivity of fish to AHR agonists, like dioxin, is far greater than what has been observed for many other vertebrates, including humans, complicating the ability to extrapolate the results of PAH toxicity studies to other species. In contrast, the early life stages of *Xenopus laevis* provide an attractive system for investigating the developmental impacts of PAH exposure because, in comparison to fish and even rodent models, the sensitivity of their AHR system to certain AHR agonists (e.g. dioxin) is greatly reduced and more closely mirrors that of humans and other amphibians (Lavine et al., 2005; Silkworth et al., 2005; Zimmermann et al., 2008). Comparing the responses to developmental exposures of single PAHs between two important model systems that vary greatly in their degree of AHR sensitivity could reveal valuable information about mechanism(s) of PAH cardiotoxicity.

Despite differences in heart morphology, the progression of heart development, including the timing of cardiac looping (by 36 hpf) and valve formation (72 hpf), is very similar for *Xenopus laevis* and zebrafish (*Danio rerio*) allowing for ready comparison of PAH-induced cardiotoxicity between species. Like zebrafish embryos, *Xenopus* embryos and larvae are transparent, allowing one to observe for abnormalities in cardiac function. As with zebrafish (*Danio rerio*), *Xenopus* embryos can rely on diffusion alone until day 7-8 of development (Burggren, 2004) which includes the entire period of heart development. Indeed, despite the presence of gross cardiac abnormalities, the lethal effects of PAH-induced cardiotoxicity in zebrafish do not manifest until day 7 when diffusion becomes limiting (Incardona et al., 2006). Our survival data support this as the case for the early life stages of *Xenopus* as well, as no increase in PAH-induced mortality was observed until day 8 of development (Figure 1).

To varying degree, exposure to all of the PAHs under investigation here disrupted the normal pacemaker activity of *Xenopus* embryos and/or larvae at some point during the period of heart development (Figure 2). Only the highest dose of FLA (25 μM) increased HR after 48 h of exposure (stage 37, during atrioventricular partitioning and immediately following cardiac looping). In contrast, by NF stage 42 (during atrial partitioning, 72 hpe), embryos exposed to PYR, PHE, or FLA at 0.25-25 μM or to BaP at 5 μM exhibited ventricular tachycardia (increased heart rate). Interestingly, these PAH-induced increases in HR were not dose-dependent, an indication that heart rates may have been at their maximum. In general, PAH-induced tachycardia persisted into stage 45 (valve formation, 96 hpe) for all but the highest doses of PHE and FLA which did not differ from controls. As development progressed, ventricular tachycardia was no longer observed with exposure to any of the PAHs (mature heart phenotype, stage 46/47,120 hpe). However, larvae exposed to FLA at 25 μM exhibited bradycardia (reduced HR).

Atrial beating rates (Figure 2) typically mirrored the PAH-induced responses observed for ventricular rates with one notable exception. At stage 46/47, atrial tachycardia was associated with FLA exposure at 2.5 and 25 μM, yet ventricular bradycardia was observed 25 μM, indicating that FLA interferes with normal cardiac conduction in a dose-dependent manner. In general, PAH exposure was associated with increased incidence of atrioventricular (AV) block, as assessed by the difference in atrial and ventricular beating rates, in a dose-dependent manner (Table 4), ranging from 19 – 45% of animals exhibiting AV block with exposure to the highest doses of PYR, PHE, and FLA. The predominant form of AV block observed was 2:1 AV block (two atrial contractions for every ventricular contraction) with some individuals in ventricular standstill for the entire 20 s period. Instances of AV block were noted with BaP exposure but to a lesser degree than for the other PAHs and no BaP-exposed animals exhibited complete block.

In contrast to our results, at 72 hpe, PHE-exposed zebrafish embryos exhibit ventricular bradycardia, not tachycardia, an effect attributed to increased incidence of AV block (Incardona et al., 2005). Although a longer exposure period (120 h) was required, we did observe a trend toward PHE-induced bradycardia. In the embryos of zebrafish, but not medaka (*Oryzias melastigma*), BaP induces severe bradycardia (Incardona et al., 2011; Mu et al., 2012), a response not observed in *Xenopus* embryos or larvae.

Studies using fish embryos have not revealed evidence of AV block or compromised cardiac conduction activity with exposures to PYR or FLA, as single compounds, as was observed here with *Xenopus.* In particular, the degree of disruption of normal pacemaker activity with FLA exposure was surprising, because, to our knowledge, as a single compound, FLA is not reported to alter pacemaker activity in fish embryos. However, AV conduction block has been previously associated with PHE exposure in fish (Incardona et al., 2004). Specifically, fish exposed to PHE as a single compound or as a component of weathered crude oil have demonstrated an array of cardiac dysfunctions including bradycardia, reduced contractility, arrhythmias, pericardial edema, and heart failure, effects that can be explained by disruption of normal cardiac conduction.

As predicted, the disruption of normal pacemaker and conduction activity observed with BaP exposure was less impactful than for the other PAHs and certainly in comparison to BaP-induced effects on the early fish heart, which are quite severe. In *Xenopus*, BaP exposure was associated with atrial and ventricular tachycardia but only at the highest dose, a lower incidence of AV block, no evidence of complete block, and, also in contrast to the other PAHs, no increase in mortality by day 8 (Figure 1). And, while BaP exposure was associated with higher incidence of developmental abnormalities in comparison to the other PAHs (Table 3), the percentage of affected animals remained low. Only 7% of BaP-exposed animals had craniofacial abnormalities and no edema was observed, in stark contrast to observations with BaP exposure in fish embryos (Huang et al., 2012; Incardona et al., 2011). Morpholino studies in fish reveal that BaP-induced developmental abnormalities and cardiotoxicity are dependent on AHR activation and CYP1A induction (Incardona et al., 2011). Because the early life stages of *Xenopus* are known to be resistant to potent AHR agonists like dioxin, a reduced impact on normal cardiac function for BaP was predicted and observed for this study.

Interestingly, the comparatively reduced impact on normal cardiac function seen with BaP exposure was accompanied by the most potent degree of AHR activation of all the PAHs, a 60-fold increase in mRNA levels for cyp1a7, the more BaP-sensitive of the paralogs (Figure 3). While PHE is not reported to stimulate AHR activity in fish, PHE was associated with a 15-fold increase in *cyp1a6* expression in our study. Furthermore, exposure to PYR, purportedly a weak AHR agonist in fish (Incardona et al., 2006), did not enhance the expression of either *cyp1a* gene. The most surprising finding, however, was that FLA, an inhibitor of CYP1A in zebrafish, was found to act as an AHR agonist (25 μM) in *Xenopus* embryos, as evidenced by a 35-fold increase in *cyp1a7* expression. The finding of cardiotoxicity and increased *cyp1a* expression with exposure to FLA in *Xenopus* was unexpected. Previous studies, in fish, have observed cardiotoxic impacts with FLA only when combined with other PAHs, like BaP (Van Tiem and Di Giulio, 2011). We are unable to conclude from this study whether FLA-induced cardiotoxicity is related to CYP1A activity in *Xenopus.*

**Figure 3.**
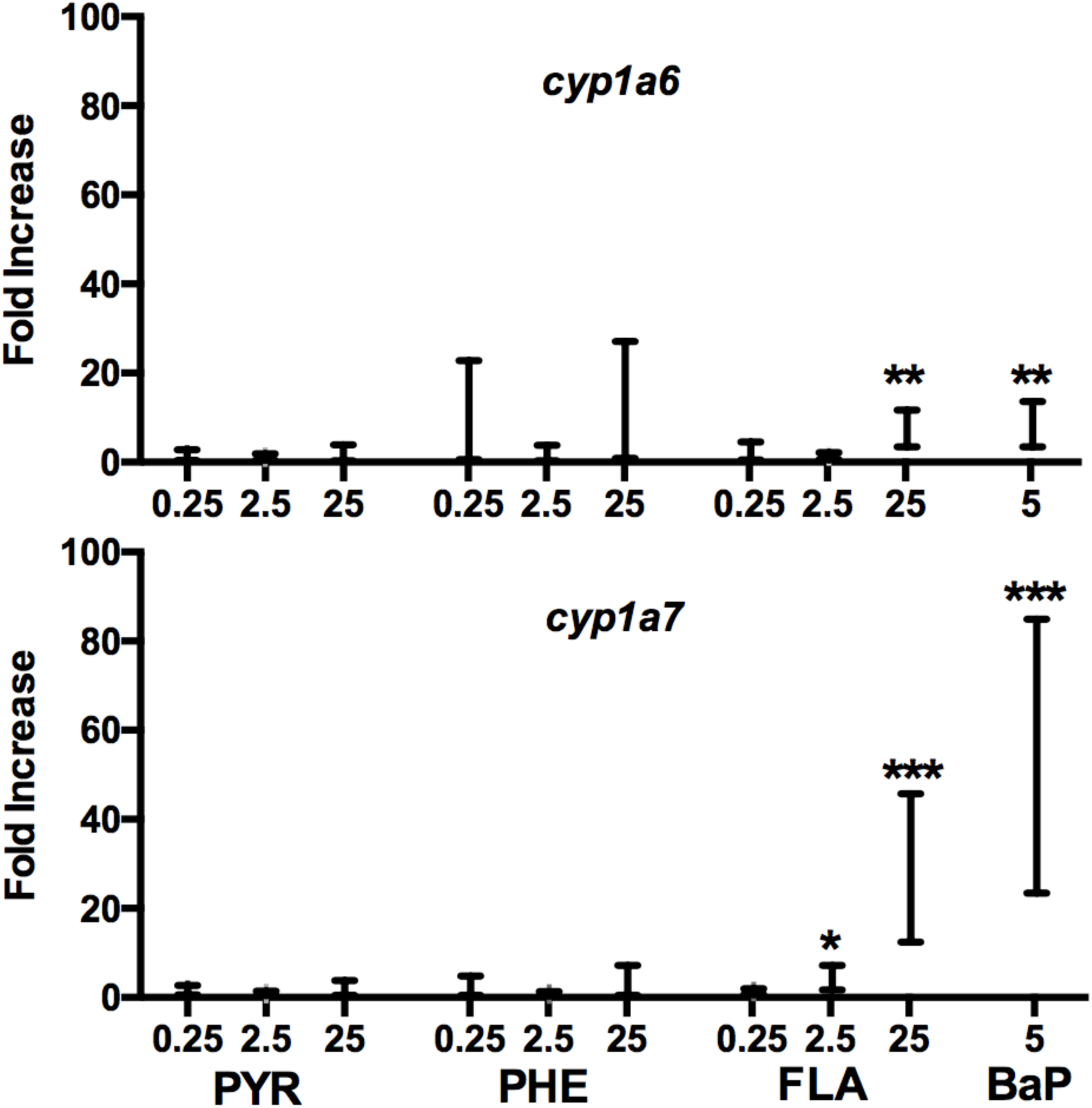
Fold change in mRNA expression, relative to vehicle control (=1), for cyp*1a6* and *cyp1a7* genes following 72 h PAH exposures. Values were calculated using the 2^-ΔΔCT^ method and are expressed as a range (incorporating the standard deviation). Each sample represents a pool of 7-9 embryos (NF stage 42) (n= 4-6 samples/treatment; 3 batches). Asterisks indicate a difference from the vehicle control (^⋆^ p <0.05, ^⋆⋆^ p <0.01, ^⋆⋆⋆^ p <0.001) PYR = pyrene, PHE = phenanthrene, FLA = fluoranthene (at 0.25, 2.5 or 25 μM), and BaP = benzo(a)pyrene (at 5 μM).

In *Xenopus*, there was no apparent correlation between cardiotoxicity and *cyp1a* mRNA levels. BaP, arguably the least impactful of normal cardiac function, was the most potent AHR agonist. In addition, significant cardiac impacts were observed with exposure to any of the three other PAHs, but only the highest dose of FLA (25 μM) stimulated *cyp1a* mRNA expression to any measurable degree. And, while these findings do not constitute proof, they do not support the idea that the cardiotoxic impacts of PHE or PYR are likely to be AHR-mediated in *Xenopus.* There remains the possibility that the AHR plays a mechanistic role in FLA-induced effects on cardiac function in *Xenopus*, but it is important to recognize that *cyp1a* mRNA levels are normally extremely low in early life stages of *Xenopus* (Zimmermann et al., 2008), making it difficult to predict whether an increase of 35-fold, as seen with 25 μM FLA, is of any physiological relevance. Based on the results reported here, alternative mechanisms should be explored.

The cardiotoxicity of weathered crude oil, enriched with PHE and other three-ringed PAHs, was investigated using electrophysiological and Ca^2+^ -imaging techniques on isolated cardiomyocytes obtained from juvenile tuna hearts (Brette et al., 2014). The degree of cardiotoxicity of different oil samples correlated with their concentrations of three-ringed PAHs including PHE. Crude oil exposure was associated with a significant delay in repolarization, lengthening the duration of the action potential (APD), an effect attributed to a decrease in the rectifier potassium (I_Kr_) current due to physical blockade of the ion channel. Decrease in the calcium current (I_Ca_) and in calcium cycling, disrupting normal excitation-contraction coupling, was also observed. Since we observed similar cardiac abnormalities with PHE exposure in *Xenopus*, it is likely that these same mechanisms are at play. The observation that, in *Xenopus*, FLA exerts cardiotoxic impacts similar to those observed with PHE suggests, that FLA may also physically disrupt ion channels critical to normal cardiac function.

In summary, these results confirm previous studies demonstrating compound-specific effects on normal pacemaker and conduction activity with early PAH exposure. They also reveal species-specific differences between the PAH-induced cardiotoxic responses of fish and frog, an important contribution since nearly all current studies focus on fish models. Finally, our findings have implications for the normal development of the cardiovascular systems of PAH-exposed humans and wild amphibians.

## Funding

Research reported in this publication was supported by an Institutional Development Award (IDeA) from the National Institute of General Medical Sciences of the National Institutes of Health under grant number P20GM103506.

## Acknowledgements

The authors would like to thank Dr. Leslie Henderson and Donna Porter of the Geisel School of Medicine at Dartmouth College and Brian Moore at Keene State College for their support of the study. The research efforts of several previous Keene State College students were instrumental to our ability to complete the experiments described here. We acknowledge the contributions of Jeffrey Hall, Krist Hausken, Jade Halsey, Elizabeth Richards, Christin Gaudette, Deena Snoke, Kaitlin Rosato, and Luke Hebert. Special recognition is given to Beth A. Freeman for scoring embryos for developmental abnormalities. The authors have no conflict of interest with regard to any content included in this article.

